# Genetic enhancement of inbred lines through pre-breeding in maize

**DOI:** 10.1101/2025.07.23.666412

**Authors:** O Srikanth, B Jyothi, MV Nagesh Kumar, Chikkappa G Karjagi, Vanishree, N Sunil

## Abstract

The improvement in the genetic yield potential of maize has led to the development of an extremely large number of maize cultivars across almost all maize-growing countries in the world. In the process, the genetic diversity in the modern maize cultivars has gradually narrowed down. To improve the genetic diversity, there is a need to undertake pre-breeding through wide-hybridization between inbred lines and their wild species, namely *Zea mays* ssp. *Parvigluimis* (WS-5) and *Zea luxurians* (WS-1). In the present study, 69 elite inbred lines were crossed with two wild relatives of maize, *Z. mays* ssp. *Parviglumis* and *Z. luxurians*. The resulting hybrids of wide hybridization showed typical wild species characters like tillering, prolificacy on main tiller and side tillers, and ear characters. This shows the introgression of characters from wild species. Combining ability analysis revealed the presence of a significant amount of variability among parents and crosses for most traits under study. *Z. luxurians* was a good general combiner for all the traits except tassel length, anthesis silking interval, and days to 75% dry husk based on the GCA effects. Whereas lines, IC0621049 and IC0621565, were good general combiners for all the traits except days to 75% dry husk. Among the crosses, highly significant SCA effects in the desired direction were shown by PFSR-3 × WS-5, EC618215 × WS-1, and IC213122 × WS-1 for plant height, ear diameter, no. of kernel rows per ear, no. of kernels per row, ear weight, and grain yield per ear. Molecular analysis also confirmed the genuine nature of hybrids. Cluster analysis classified all the parents involved in the crosses into two groups, with most of the inbred lines grouping with WS-5, revealing that most of the inbred lines were closely related to WS-5 as compared to WS-1. This may be the reason for the development of more successful crosses with WS-5 as compared to WS-1.

## INTRODUCTION

Maize (*Zea mays* L.) is the 3^rd^ most important cereal crop; it is grown across the tropical, subtropical, and temperate climatic regions. It became important because of its diverse uses in the form of food, animal feed, biofuel, and various industrial processes (Cassidy *et al*., 2013). Domestication followed by selection for desired traits narrowed down the genetic diversity in modern maize cultivars in comparison to their wild relatives (Vigouroux *et al*., 2005). Decrease in genetic diversity leads to genetic vulnerability, which ultimately reduces production due to various biotic and abiotic stresses (Rosegrant *et al*., 2009). In addition, climate change predictions reported that agricultural productivity would be significantly harmed, and many regions wouldn’t be able to make the improvements required for long-term food security (Cairns *et al*., 2012).

To counteract these challenges, plant breeders will need all the genetic diversity that they can get. Landraces and heirloom cultivars, which are still grown by farmers all over the world, have some of this diversity but have low yields, which is a major bottleneck. So, to introduce this diversity into present inbred lines, a pre-breeding programme must be employed for genetic enhancement of inbred lines (Dempewolf *et al*., 2014). Wild crop relatives have played a key role in understanding plant genomes and improving cultivated crops by contributing valuable genetic traits, leading to major advances in both research and breeding. (Mammadov *et al*.,2018).

Keeping this in consideration, in this study, maize inbred lines were crossed with their wild relatives, such as *Zea mays* ssp. *Parvigluimis* (WS-5) and *Zea luxurians* (WS-1) in L × T fashion. Resulting crosses were evaluated to know about the expression of traits from wild species in the hybrids and for combining ability. In addition, a molecular study was done to know the true type of hybrids.

## MATERIALS AND METHODS

The present research includes crossing between 69 inbred lines (obtained from IIMR, Ludhiana) and two wild species such as WS-5 and WS-1, in *Kharif*, 2022, resulting in 58 successful crosses. Among them, 46 lines were involved in crossing with WS-5, and 12 lines were involved in crossing with WS-1; this creates an imbalance of crossings. Of the 58 inbred lines, eight inbred lines were common female parents with both wild species that developed hybrids. Hybrids were evaluated in *Rabi*, 2022-23 at Winter Nursery Centre, Rajendranagar, Hyderabad. In parents and hybrids, data were recorded on three randomly selected plants for traits like plant height, ear length, ear diameter, number of kernel rows per ear, number of kernels per row, ear weight, and grain yield in each replication. Flowering and maturity traits, *i*.*e*., days to 50% tasseling, days to 50% silking, anthesis silking interval, and days to 75% dry husk. In-addition for wild species data was recorded in traits like days to anthesis, days to silking, number of tillers per plant, number of tassels per plant, number of silks per plant, number of nodes per plant, internodal length, leaf length and leaf width and in hybrids data was recorded for number of tillers and number of silks.

### DNA Extraction

Molecular analysis was carried out using DNA isolated from leaves collected from parents and the resulting hybrids at 30 days 30-day-old seedling stage, and DNA was isolated using the CTAB (Cetyltrimethyl ammonium bromide) method. The quality of DNA was assessed using a Nanodrop 1000 spectrophotometer. All the samples were diluted to 10 ng/µl after measurement and utilised in the polymerase chain reaction.

### PCR amplification

Twelve markers were used to determine the purity of hybrids, and the detailed information about the markers can be found at http://maize.gdb. The PCR reaction mixture was made up to 10 µl (2.0 μl DNA, 0.5 μl Forward primer, 0.5 μl Reverse primer, 4.0 µl Premix (Takara), and 3.0 µl sterile distilled water). PCR reaction was performed in Biosystems™Veriti™ 96 well thermal cycler and PCR conditions were initial denaturation at 95° C for 5 min and followed by 30 cycles of denaturation at 95° C for 45 seconds, annealing at 54-58 degree C for 45 seconds and extension at 72° C for 1 minute and final extension at 72° C for 10 minutes. The amplified PCR products were resolved on a 3% Agarose gel by using a gel electrophoresis unit and were visualized under the UV transilluminator and documented using a gel documentation system (BIO-RAD, BioRad Laboratories, India).

### Data analysis

Analysis for combining ability was done using Henderson’s (1952) analysis method L×T package developed by CIMMYT, Mexico, which was developed for imbalanced crossings. For molecular analysis binary data matrix (score ‘1’ and ‘0’) of the alleles of the SSR marker obtained from parents and hybrids was subjected to grouping using Darwin 6.0.021 software (Perrier and Jacquemoud, 2006).

## RESULTS AND DISCUSSION

### Morphological characterisation of wild species

Wild species showed typical characteristics that were absent in inbred lines. Data were recorded for wild species characteristics, and the mean values for the traits recorded were represented in Table 1. The traits revealed that the wild species showed protogyny, whereas present maize cultivars are protandrous, and the wild species had tillering, prolificacy on main tiller and side tillers, and multiple tassels per plant, which were absent in the inbred lines. In wild species, multiple tassels had emerged from side branches, and silk/ear emerged from nodes. On average, every node bears a silk. Wild species had more nodes and had high internodal length as compared to present maize cultivars. Wild species had very low ear length and the number of kernel rows per ear as compared to modern maize cultivars, depicting the influence of artificial selection on maize. A similar kind of result was reported by Adhikari *et al*. (2019).

**Table 1.**
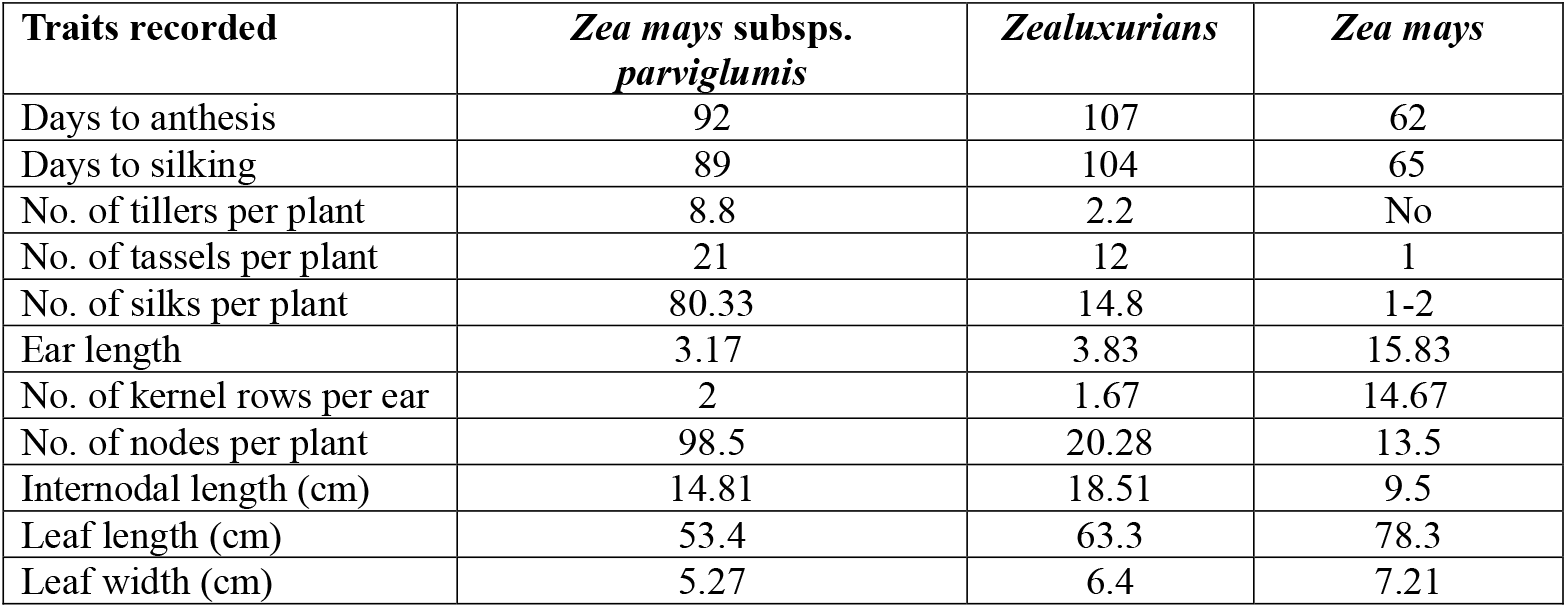
Mean values for various traits recorded in wild species.

### Morphological characterization of hybrids

In hybrids, typical wild species characteristics like tillering and prolificacy on main tillers and side tillers were observed, which were depicted in Table 2. On average, no. of tillers ranged from 1 to 6.6, and for no. of silks from 1 to 6.4. In F_1_’s, IC0621166 × WS-5 (6.6) showed highest number of tillers then by IC0621634 × WS-5 (6.5), then IC212875 × WS-5 (6.3) followed by IC420922 × WS-1 and IC522300 × WS-1(6.2). Whereas for silking habit F_1_, IC0612832 × WS-5 (6.5) showed highest number of silks then by IC0612832 × WS-5 (6.4) followed by MIL 6-2 × WS-5 and IC0621166 × WS-5 (5.5). This shows the dominant expression of an introgressed trait of wild species in the F_1_ hybrids. Figures 1 and 2 represent prolificacy on the main tiller and side tillers and tillering habitat in hybrids, respectively.

**Table 2.** Mean values for No. of tillers and silks observed in hybrids.

| Hybrid | No. of tillers | No. of Silks |
| --- | --- | --- |
| BJIM-08-27× WS5 | 4.6 | 3.3 |
| BML-7× WS-5 | 2.5 | 3 |
| EC0758189× WS-5 | 6 | 2 |
| EC618215× WS-1 | 5.3 | 4.5 |
| EC618215× WS-5 | 1 | 1 |
| HKI-163× WS-1 | 4.3 | 4 |
| HKI-163× WS-5 | 3 | 1 |
| IC0597385× WS-5 | 1 | 2 |
| IC0598546× WS-5 | 3.7 | 4 |
| IC0612729× WS-5 | 3 | 4 |
| IC0612770× WS-1 | 4.6 | 2 |
| IC0612770× WS-5 | 8.3 | 1 |
| IC0612821× WS-1 | 4.6 | 3 |
| IC0612832× WS-5 | 5.4 | 6.4 |
| IC0612864× WS-5 | 4.2 | 5 |
| IC0616502× WS-5 | 5 | 2 |
| IC0620929× WS-5 | 2.3 | 2.3 |
| IC0620969× WS-5 | 4.25 | 3.3 |
| IC0620978× WS-5 | 5.2 | 5 |
| IC0621049× WS-1 | 1.5 | 1 |
| IC0621061× WS-5 | 4.5 | 2.7 |
| IC0621159× WS-5 | 4.3 | 4.6 |
| IC0621166× WS-5 | 6.6 | 5.5 |
| IC0621168× WS-5 | 5.5 | 4 |
| IC0621430× WS-5 | 5.9 | 1 |
| IC0621565× WS-5 | 1.8 | 1 |
| IC0621578× WS-5 | 3.5 | 2 |
| IC0621634× WS-5 | 6.5 | 4 |
| IC212838× WS-5 | 3 | 3 |
| IC212871× WS-5 | 2 | 4 |
| IC212875× WS-1 | 4 | 7 |
| IC212875× WS-5 | 6.3 | 1 |
| IC212876× WS-5 | 4.8 | 3 |
| IC213004× WS-5 | 5 | 4 |
| IC213007× WS-5 | 1.2 | 1 |
| IC213035× WS-5 | 2 | 5.2 |
| IC213053× WS-5 | 2.5 | 4 |
| IC213068× WS-1 | 2 | 1 |
| IC213090× WS-5 | 5 | 2.5 |
| IC213120× WS-5 | 3.4 | 1.5 |
| IC213122× WS-1 | 2 | 1 |
| IC213122× WS-5 | 4.6 | 3.4 |
| IC213132× WS-1 | 2.2 | 3.3 |
| IC213132× WS-5 | 3.8 | 1.8 |
| IC254061× WS-5 | 5 | 4.2 |
| IC254062× WS-5 | 5.2 | 3.8 |
| IC420922× WS-1 | 6.2 | 4.5 |
| IC522300× WS-1 | 6.2 | 8 |
| IC522300× WS-5 | 1.5 | 1 |
| IC563961× WS-5 | 2 | 1 |
| MIL6-1× WS-5 | 1.25 | 1 |
| MIL6-19× WS-5 | 1 | 1 |
| MIL6-2× WS-5 | 3 | 5.5 |
| MIL6-26× WS-5 | 6.1 | 5 |
| MIL6-P6× WS-5 | 2.5 | 3.4 |
| MIL6-P6M× WS-5 | 1.5 | 1 |
| PFSR-3× WS-1 | 4.2 | 5 |
| PFSR-3× WS-5 | 3 | 1 |

**Figure 1.**
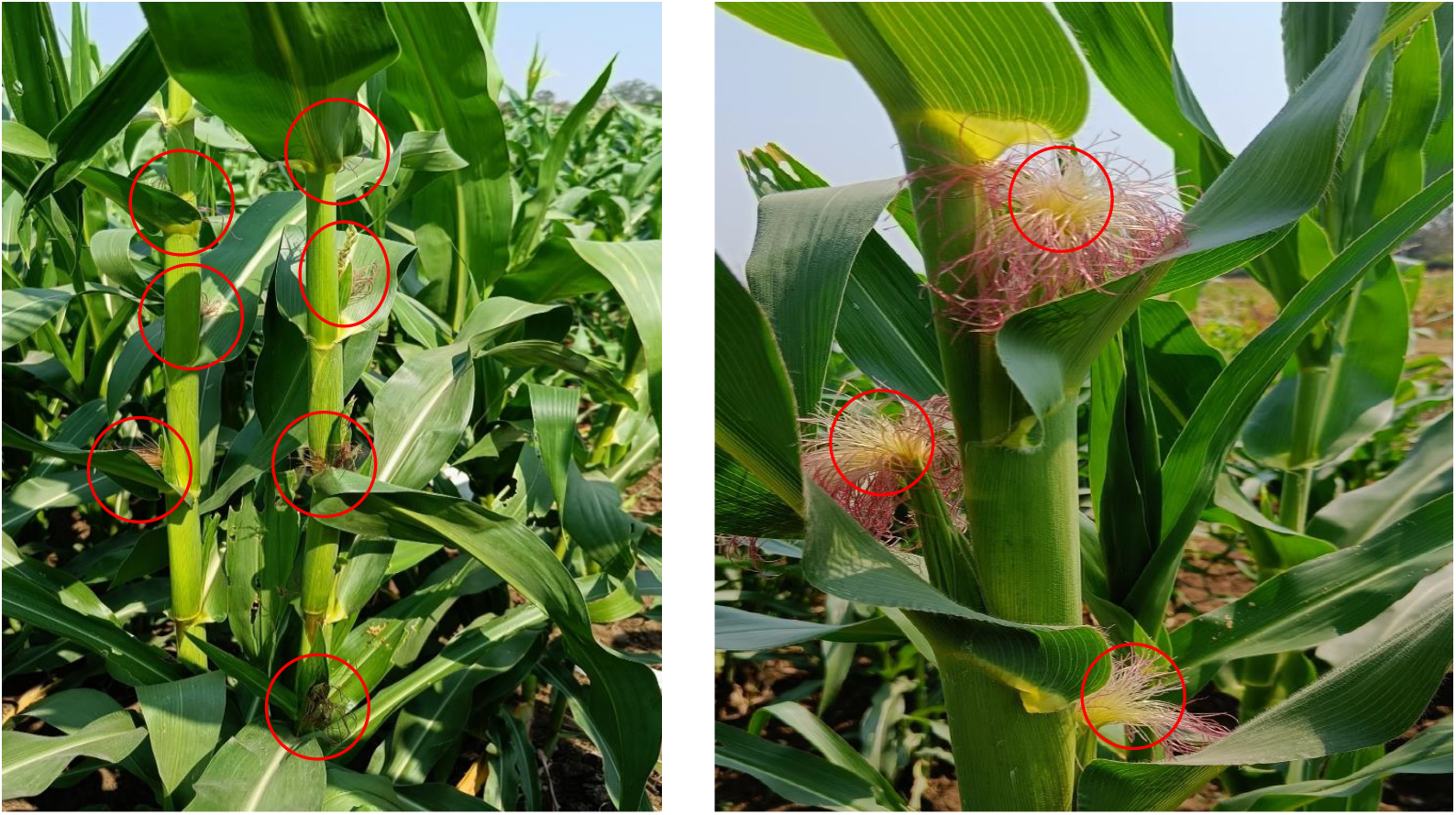
Prolificacy on the main tiller and side tillers habitat observed in hybrids.

**Figure 2.**
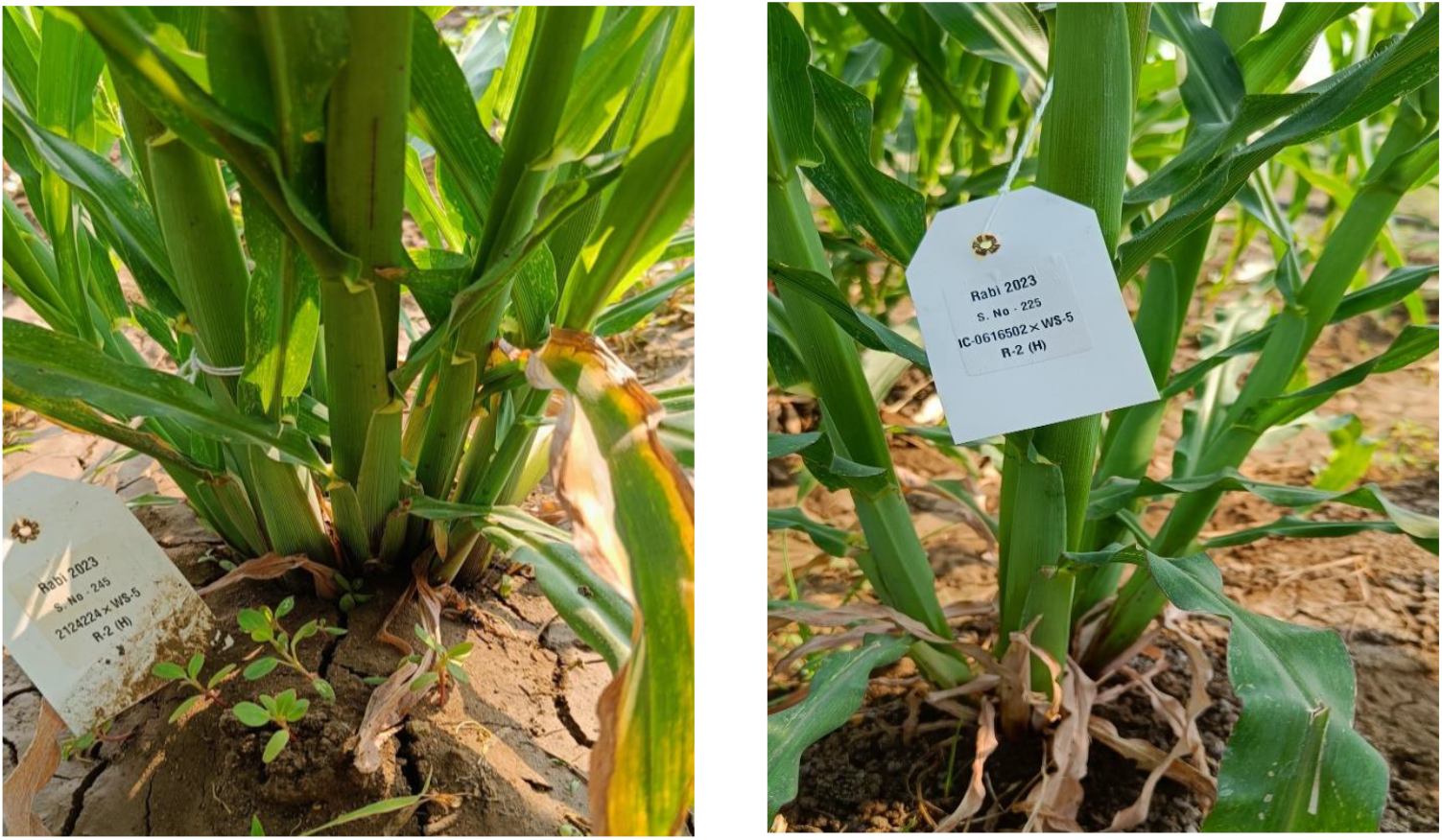
Tillering habitat observed in hybrids.

### Combining ability in maize

Analysis of variance (Table 3) for combining ability revealed the presence of significant differences in replications for traits, namely plant height, tassel length, days to 50% tasseling, days to 50% silking, anthesis silking interval, and days to 75% dry husk. Whereas in treatments, all the characters under study showed significant differences, indicating that there was a presence of variation among the genotypes. Among the parents, lines, i.e., inbred lines, showed significant differences for traits, namely, days to 50% tasseling, days to 50% silking, and days to 75% dry husk. Whereas testers didn’t show any significant differences. In the case of hybrids, all traits under investigation showed a significant difference. Even though the variation was non-significant for the traits in testers, crosses showed significant differences for all the traits under study. This could be due to complementary gene action of alleles at different loci, resulting in overdominance in either positive or negative directions or both directions.

**Table 3.**
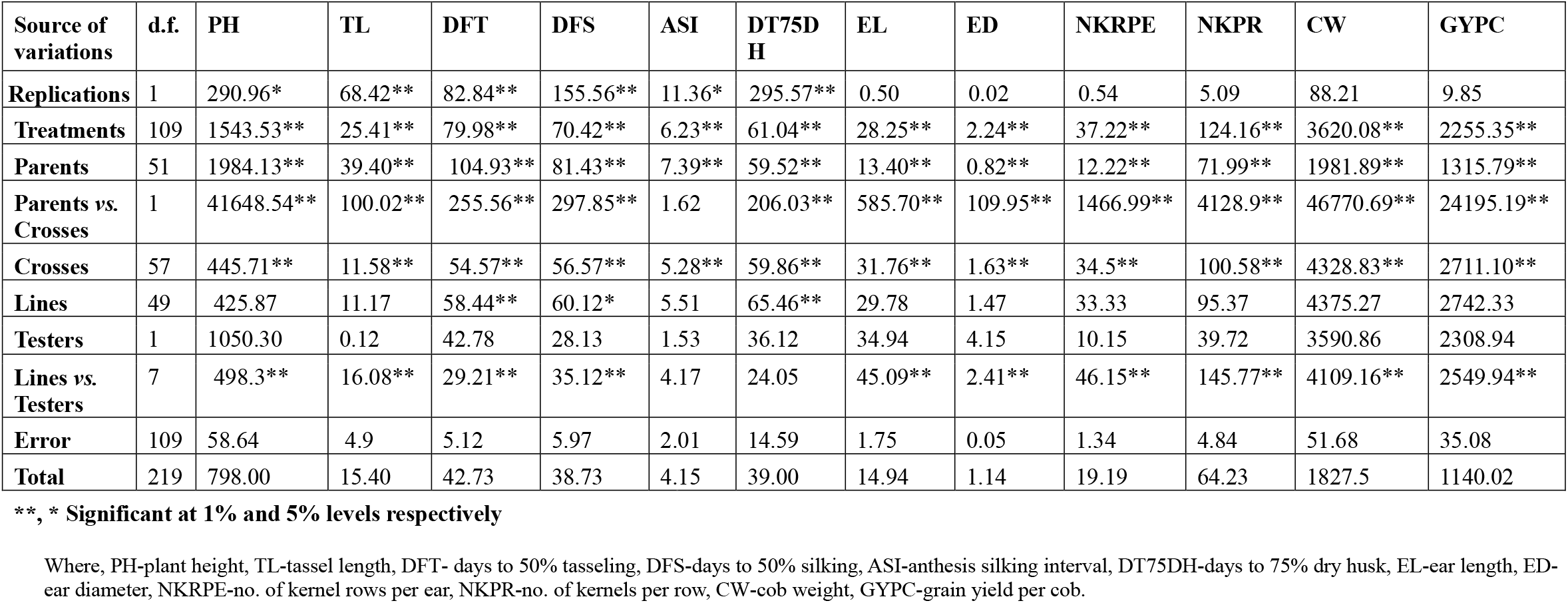
Analysis of variances for combining ability for various traits under investigation.

| Source of variations | d.f. | PH | TL | DFT | DFS | ASI | DT75D<br>H | EL | ED | NKRPE | NKPR | CW | GYPC |
| --- | --- | --- | --- | --- | --- | --- | --- | --- | --- | --- | --- | --- | --- |
| **Replications** | 1 | 290.96* | 68.42** | 82.84** | 155.56** | 11.36* | 295.57** | 0.50 | 0.02 | 0.54 | 5.09 | 88.21 | 9.85 |
| **Treatments** | 109 | 1543.53** | 25.41** | 79.98** | 70.42** | 6.23** | 61.04** | 28.25** | 2.24** | 37.22** | 124.16** | 3620.08** | 2255.35** |
| **Parents** | 51 | 1984.13** | 39.40** | 104.93** | 81.43** | 7.39** | 59.52** | 13.40** | 0.82** | 12.22** | 71.99** | 1981.89** | 1315.79** |
| **Parents vs. Crosses** | 1 | 41648.54** | 100.02** | 255.56** | 297.85** | 1.62 | 206.03** | 585.70** | 109.95** | 1466.99** | 4128.9** | 46770.69** | 24195.19** |
| **Crosses** | 57 | 445.71** | 11.58** | 54.57** | 56.57** | 5.28** | 59.86** | 31.76** | 1.63** | 34.5** | 100.58** | 4328.83** | 2711.10** |
| **Lines** | 49 | 425.87 | 11.17 | 58.44** | 60.12* | 5.51 | 65.46** | 29.78 | 1.47 | 33.33 | 95.37 | 4375.27 | 2742.33 |
| **Testers** | 1 | 1050.30 | 0.12 | 42.78 | 28.13 | 1.53 | 36.12 | 34.94 | 4.15 | 10.15 | 39.72 | 3590.86 | 2308.94 |
| **Lines vs. Testers** | 7 | 498.3** | 16.08** | 29.21** | 35.12** | 4.17 | 24.05 | 45.09** | 2.41** | 46.15** | 145.77** | 4109.16** | 2549.94** |
| **Error** | 109 | 58.64 | 4.9 | 5.12 | 5.97 | 2.01 | 14.59 | 1.75 | 0.05 | 1.34 | 4.84 | 51.68 | 35.08 |
| **Total** | 219 | 798.00 | 15.40 | 42.73 | 38.73 | 4.15 | 39.00 | 14.94 | 1.14 | 19.19 | 64.23 | 1827.5 | 1140.02 |

**\*\* , \* Significant at 1% and 5% levels respectively**
Where, PH-plant height, TL-tassel length, DFT- days to 50% tasseling, DFS-days to 50% silking, ASI-anthesis silking interval, DT75DH-days to 75% dry husk, EL-ear length, ED- ear diameter, NKRPE-no. of kernel rows per ear, NKPR-no. of kernels per row, CW-cob weight, GYPC-grain yield per cob.

The trait-wise estimates of the effects of GCA and SCA were presented in Tables 4 and 5, respectively. The GCA effects revealed that none of the lines or testers were good general combiners for all twelve quantitative characters studied. However, among the testers, WS-1 was a good general combiner for all the traits except tassel length, anthesis silking interval, and days to 75% dry husk. Among lines EC618215, IC0621049, IC0621061, IC0621565, IC522300, and IC563961 were good general combiners for plant height, tassel length, days to 50% tasseling, days to 50% silking, and anthesis silking interval. And lines MIL6-P6M, MIL6-26, MIL6-19, MIL6-1, IC213007, IC0621565, and IC0621049 were good general combiners for ear length, ear diameter, no. of kernel rows per ear, no. of kernels per row, ear weight, and grain yield per ear.

**Table 4.**
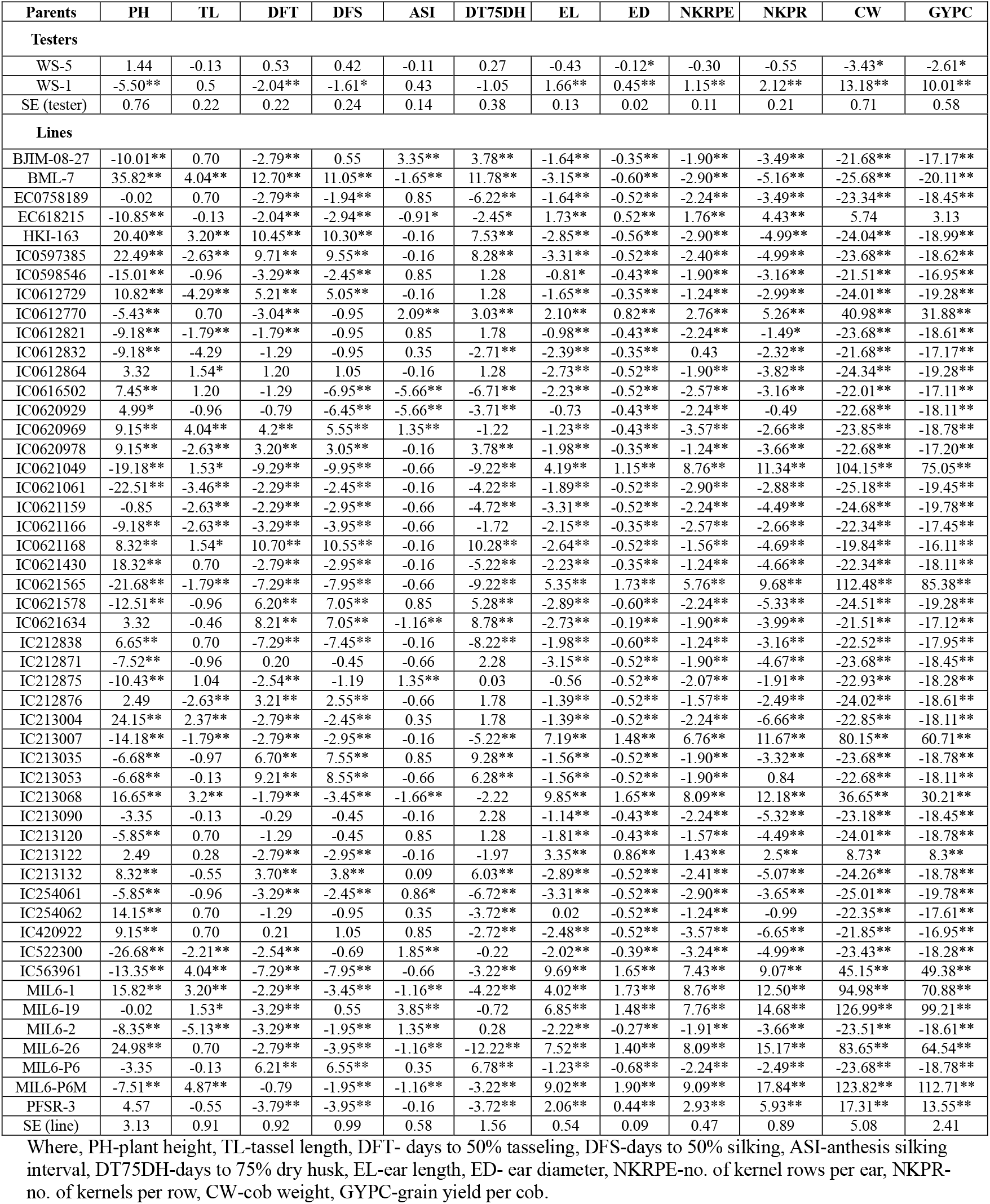
Estimates of General Combining Ability (GCA) effects of lines and testers for various traits under investigation.

**Table 5.** Estimates of Specific Combining Ability (SCA) effects for crosses of various traits under investigation.

| **Crosses** | **PH** | **TL** | **DFT** | **DFS** | **ASI** | **DT75DH** | **EL** | **ED** | **NKRPE** | **NKPR** | **CW** | **GYPC** |
| --- | --- | --- | --- | --- | --- | --- | --- | --- | --- | --- | --- | --- |
| BJIM-08-27× WS5 | -1.44 | 0.13 | -0.53 | -0.42 | 0.11 | -0.27 | 0.43 | 0.11 | 0.30 | 0.55 | 3.44 | 2.61 |
| BML-7× WS-5 | -1.44 | 0.13 | -0.53 | -0.42 | 0.11 | -0.27 | 0.43 | 0.11 | 0.30 | 0.55 | 3.44 | 2.61 |
| EC0758189× WS-5 | -1.44 | 0.13 | -0.53 | -0.42 | 0.11 | -0.27 | 0.43 | 0.11 | 0.30 | 0.55 | 3.44 | 2.61 |
| EC618215× WS-1 | -8.67* | 2.84** | 1.79 | 1.16 | -0.68 | -2.20 | 2.80** | 0.59** | 2.51** | 5.79** | 17.06** | 12.57** |
| EC618215× WS-5 | 12.73** | -3.20** | -0.28 | 0.08 | 0.36 | 2.98 | -4.03** | -0.93** | -3.36** | -7.36** | -26.81** | -19.9** |
| HKI-163× WS-1 | -0.75 | -1.33 | 1.79 | 1.87 | 0.07 | 2.78 | -1.84** | -0.41** | -1.48** | -3.45** | -12.51** | -9.22** |
| HKI-163× WS-5 | 4.86 | 0.96 | -0.28 | -0.67 | -0.39 | -2.02 | 0.61 | 0.07 | 0.63 | 1.89 | 2.77 | 1.82 |
| IC0597385× WS-5 | -1.44 | 0.13 | -0.53 | -0.42 | 0.11 | -0.27 | 0.43 | 0.12 | 0.30 | 0.55 | -13.78** | -2.61 |
| IC0598546× WS-5 | -1.44 | 0.13 | -0.53 | -0.42 | 0.11 | -0.27 | 0.43 | 0.12 | 0.30 | 0.55 | 3.44 | 2.61 |
| IC0612729× WS-5 | -1.44 | 0.13 | -0.53 | -0.42 | 0.11 | -0.27 | 0.43 | 0.12 | 0.30 | 0.55 | 3.44 | 2.61 |
| IC0612770× WS-1 | 0.92 | 0.33 | 2.79** | 3.12** | 0.32 | -2.20 | 2.26** | 0.80** | 3.18** | 6.29** | 51.82** | 41.51** |
| IC0612770× WS-5 | 3.15 | -0.70 | -1.28 | -1.92 | -0.64 | 2.98 | -3.48** | -1.13** | -4.03** | -7.86** | -61.56** | -48.55** |
| IC0612821× WS-1 | 5.5 | -0.49 | 2.04 | 1.61 | -0.43 | 1.05 | -1.66** | -0.44** | -1.15 | -2.12 | -13.17** | -10.01** |
| IC0612832× WS-5 | -1.44 | 0.13 | -0.53 | -0.42 | 0.11 | -0.27 | 0.43 | 0.12 | 0.30 | 0.55 | 3.44 | 2.61 |
| IC0612864× WS-5 | -1.44 | 0.13 | -0.53 | -0.42 | 0.11 | -0.27 | 0.43 | 0.12 | 0.30 | 0.55 | 3.44 | 2.61 |
| IC0616502× WS-5 | -1.44 | 0.13 | -0.53 | -0.42 | 0.11 | -0.27 | 0.43 | 0.12 | 0.30 | 0.55 | 3.44 | 2.61 |
| IC0620929× WS-5 | -1.44 | 0.13 | -0.53 | -0.42 | 0.11 | -0.27 | 0.43 | 0.12 | 0.30 | 0.55 | 3.44 | 2.61 |
| IC0620969× WS-5 | -1.44 | 0.13 | -0.53 | -0.42 | 0.11 | -0.27 | 0.43 | 0.12 | 0.30 | 0.55 | 3.44 | 2.61 |
| IC0620978× WS-5 | -1.44 | 0.13 | -0.53 | -0.42 | 0.11 | -0.27 | 0.43 | 0.12 | 0.30 | 0.55 | 3.44 | 2.61 |
| IC0621049× WS-1 | 5.5 | -0.49 | 2.04 | 1.61 | -0.43 | 1.05 | -1.66** | -0.45** | -1.15 | -2.12 | -13.18** | -10.01** |
| IC0621061× WS-5 | -1.44 | 0.13 | -0.53 | -0.42 | 0.11 | -0.27 | 0.43 | 0.11 | 0.30 | 0.55 | 3.44 | 2.61 |
| IC0621159× WS-5 | -1.44 | 0.13 | -0.53 | -0.42 | 0.11 | -0.27 | 0.43 | 0.11 | 0.30 | 0.55 | 3.44 | 2.61 |
| IC0621166× WS-5 | -1.44 | 0.13 | -0.53 | -0.42 | 0.11 | -0.27 | 0.43 | 0.11 | 0.30 | 0.55 | 3.44 | 2.61 |
| IC0621168× WS-5 | -1.44 | 0.13 | -0.53 | -0.42 | 0.11 | -0.27 | 0.43 | 0.11 | 0.30 | 0.55 | 3.44 | 2.61 |
| IC0621430× WS-5 | -1.44 | 0.13 | -0.53 | -0.42 | 0.11 | -0.27 | 0.43 | 0.11 | 0.30 | 0.55 | 3.44 | 2.61 |
| IC0621565× WS-5 | -1.44 | 0.13 | -0.53 | -0.42 | 0.11 | -0.27 | 0.43 | 0.11 | 0.30 | 0.55 | 3.44 | 2.61 |
| IC0621578× WS-5 | -1.44 | 0.13 | -0.53 | -0.42 | 0.11 | -0.27 | 0.43 | 0.11 | 0.30 | 0.55 | 3.44 | 2.61 |
| IC0621634× WS-5 | -1.44 | 0.13 | -0.53 | -0.42 | 0.11 | -0.27 | 0.43 | 0.11 | 0.30 | 0.55 | 3.44 | 2.61 |
| IC212838× WS-5 | -1.44 | 0.13 | -0.53 | -0.42 | 0.11 | -0.27 | 0.43 | 0.11 | 0.30 | 0.55 | 3.44 | 2.61 |
| IC212871× WS-5 | -1.44 | 0.13 | -0.53 | -0.42 | 0.11 | -0.27 | 0.43 | 0.11 | 0.30 | 0.55 | 3.44 | 2.61 |
| IC212875× WS-1 | -0.75 | -3.33** | 1.29 | -1.14 | -2.43** | -0.20 | -1.57** | -0.36** | -1.65** | -3.20** | -13.93** | -10.34** |
| IC212875× WS-5 | 4.82 | 2.95** | 0.21 | 2.33 | 2.11** | 0.97 | 0.35 | 0.03 | 0.80 | 1.63 | 4.18 | 2.95 |
| IC212876× WS-5 | -1.44 | 0.13 | -0.53 | -0.42 | 0.11 | -0.27 | 0.43 | 0.12 | 0.30 | 0.55 | 3.44 | 2.61 |
| IC213004× WS-5 | -1.44 | 0.13 | -0.53 | -0.42 | 0.11 | -0.27 | 0.43 | 0.12 | 0.30 | 0.55 | 3.44 | 2.61 |
| IC213007× WS-5 | -1.44 | 0.13 | -0.53 | -0.42 | 0.11 | -0.27 | 0.43 | 0.12 | 0.30 | 0.55 | 3.44 | 2.61 |
| IC213035× WS-5 | -1.44 | 0.13 | -0.53 | -0.42 | 0.11 | -0.27 | 0.43 | 0.12 | 0.30 | 0.55 | 3.44 | 2.61 |
| IC213053× WS-5 | -1.44 | 0.13 | -0.53 | -0.42 | 0.11 | -0.27 | 0.43 | 0.12 | 0.30 | 0.55 | 3.44 | 2.61 |
| IC213068× WS-1 | 5.5 | -0.49 | 2.04 | 1.66 | -0.43 | -1.05 | -1.66** | -0.45** | -1.15 | -2.12 | 13.18** | -10.14** |
| IC213090× WS-5 | -1.44 | 0.13 | -0.53 | -0.42 | 0.11 | -0.27 | 0.43 | 0.12 | 0.30 | 0.55 | 3.43 | 2.92 |
| IC213120× WS-5 | -1.44 | 0.13 | -0.53 | -0.42 | 0.11 | -0.27 | 0.43 | 0.12 | 0.30 | 0.55 | 3.44 | 2.61 |
| IC213122× WS-1 | -7.83* | -0.08 | -1.96 | -1.89 | 0.07 | -1.70 | 4.34 | 0.84** | 2.85** | 4.88** | 20.57** | 18.23** |
| IC213122× WS-5 | 11.9** | -0.29 | 3.47** | 3.08** | -0.39 | 2.47 | -5.57** | -1.18** | -3.7** | -6.45** | -30.31** | -25.63** |
| IC213132× WS-1 | -3.67 | -1.75 | -4.46** | -4.64** | -0.18 | -2.20 | -2.24** | -0.45** | -1.98** | -3.70** | -14.76** | -11.34** |
| IC213132× WS-5 | 7.73 | 1.38 | 5.96** | 5.83** | -0.14 | 2.97 | 1.01 | 0.12 | 1.13 | 2.13 | 5.02 | 3.95 |
| IC254061× WS-5 | -1.44 | 0.13 | -0.53 | -0.42 | 0.11 | -0.27 | 0.43 | 0.12 | 0.30 | 0.55 | 3.44 | 2.61 |
| IC254062× WS-5 | -1.44 | 0.13 | -0.53 | -0.42 | 0.11 | -0.27 | 0.43 | 0.12 | 0.30 | 0.55 | 3.44 | 2.61 |
| IC420922× WS-1 | 5.50 | -0.49 | 2.04 | 1.62 | -0.43 | 1.05 | -1.66** | -0.45** | -1.15 | -2.12 | -13.17** | -10.01** |
| IC522300× WS-1 | -6.99 | -1.74 | 2.29 | 3.37** | 1.07 | 2.05 | -3.37** | -0.32** | -1.15 | -3.96** | -13.26** | -9.85** |
| IC522300× WS-5 | 11.06** | 1.38 | -0.78 | -2.17 | -1.39** | -1.27 | 2.13** | -0.01 | 0.30 | 2.39* | 3.52 | 2.45 |
| IC563961× WS-5 | -1.44 | 0.13 | -0.53 | -0.42 | 0.11 | -0.27 | 0.43 | 0.12 | 0.30 | 0.55 | 3.44 | 2.61 |
| MIL6-1× WS-5 | -1.44 | 0.13 | -0.53 | -0.42 | 0.11 | -0.27 | 0.43 | 0.12 | 0.30 | 0.55 | 3.44 | 2.61 |
| MIL6-19× WS-5 | -1.44 | 0.13 | -0.53 | -0.42 | 0.11 | -0.27 | 0.43 | 0.12 | 0.30 | 0.55 | 3.44 | 2.61 |
| MIL6-2× WS-5 | -1.44 | 0.13 | -0.53 | -0.42 | 0.11 | -0.27 | 0.43 | 0.12 | 0.30 | 0.55 | 3.44 | 2.61 |
| MIL6-26× WS-5 | -1.44 | 0.13 | -0.53 | -0.42 | 0.11 | -0.27 | 0.43 | 0.12 | 0.30 | 0.55 | 3.44 | 2.61 |
| MIL6-P6× WS-5 | -1.44 | 0.13 | -0.53 | -0.42 | 0.11 | -0.27 | 0.43 | 0.12 | 0.30 | 0.55 | 3.44 | 2.61 |
| MIL6-P6M× WS-5 | -1.44 | 0.13 | -0.53 | -0.42 | 0.11 | -0.27 | 0.43 | 0.12 | 0.30 | 0.55 | 3.44 | 2.61 |
| PFSR-3× WS-1 | 25.91** | 1.59 | 3.54** | 3.61** | 0.07 | 3.55 | -5.28** | -1.41** | -6.98** | -10.71** | -55.67** | -43.34** |
| PFSR-3× WS-5 | -21.85** | -1.96 | -2.03 | -2.42 | -0.39 | -2.77 | 4.05** | 1.08** | 6.13** | 9.14** | 45.94** | 35.95** |
| SE (crosses) | 5.42 | 1.57 | 1.60 | 1.72 | 0.58 | 2.70 | 0.93 | 0.17 | 0.82 | 1.56 | 5.08 | 4.19 |

Where, PH-plant height, TL-tassel length, DFT- days to 50% tasseling, DFS-days to 50% silking, ASI-anthesis silking interval, DT75DH-days to 75% dry husk, EL-ear length, ED- ear diameter, NKPRPE-number of kernel rows per ear, NKPR of kernels per row, CW-cob weight, GYPC-grain yield per cob.

Among the 58 crosses, significant high SCA effects in desirable direction were recorded by the crosses PFSR-3 × WS-5, EC618215 × WS-1 and IC213122 × WS-1 for plant height, ear diameter, no. of kernel rows per ear, no. of kernels per row, ear weight and grain yield per ear; IC212875 × WS-1 and EC618215× WS-5 for tassel length; IC213132 × WS-1 for days to 50% tasseling and days to 50% silking; IC212875 × WS-1 and IC522300 × WS-5 for anthesis silking interval and ear length and none of the crosses showed significant SCA effects in desired direction for Days to 75% dry husk. These results are in accordance with Akshaya *et al*. (2022), Italia *et al*. (2022).

Best cross combinations with desirable SCA effects for 12 quantitative traits, along with GCA effects of their parents, revealed that the best specific combiners were resultant of good × good, good × poor, and poor × poor general combiners. The interaction between good × good combiners in a cross in plant height, ear diameter, no. of kernel rows per ear, no. Kernels per row, ear weight, and grain yield per ear can be fixed in subsequent generations, as it was due to additive × additive interaction. The superiority of crosses involving good × poor combiners could be due to additive × dominant interaction. They may produce good segregants only if general combiners were governed by additive effects and crosses were governed by epistatic effects. Whereas superior crosses involving poor × poor general combiners may be due to overdominance. The high yield of crosses involving non-additive gene action would not be fixable.

It is also observed that females (lines) made a greater contribution to the total variance for all the studied traits compared to males (testers), and the proportion of line × tester interaction to the total variance was higher than that of testers for all the traits. Proportional contribution was represented in Figure 3. For all the traits under study, the ratio between GCA to SCA was less than unity, indicating the prevalence of non-additive gene action in the inheritance of these traits.

**Figure 3.**
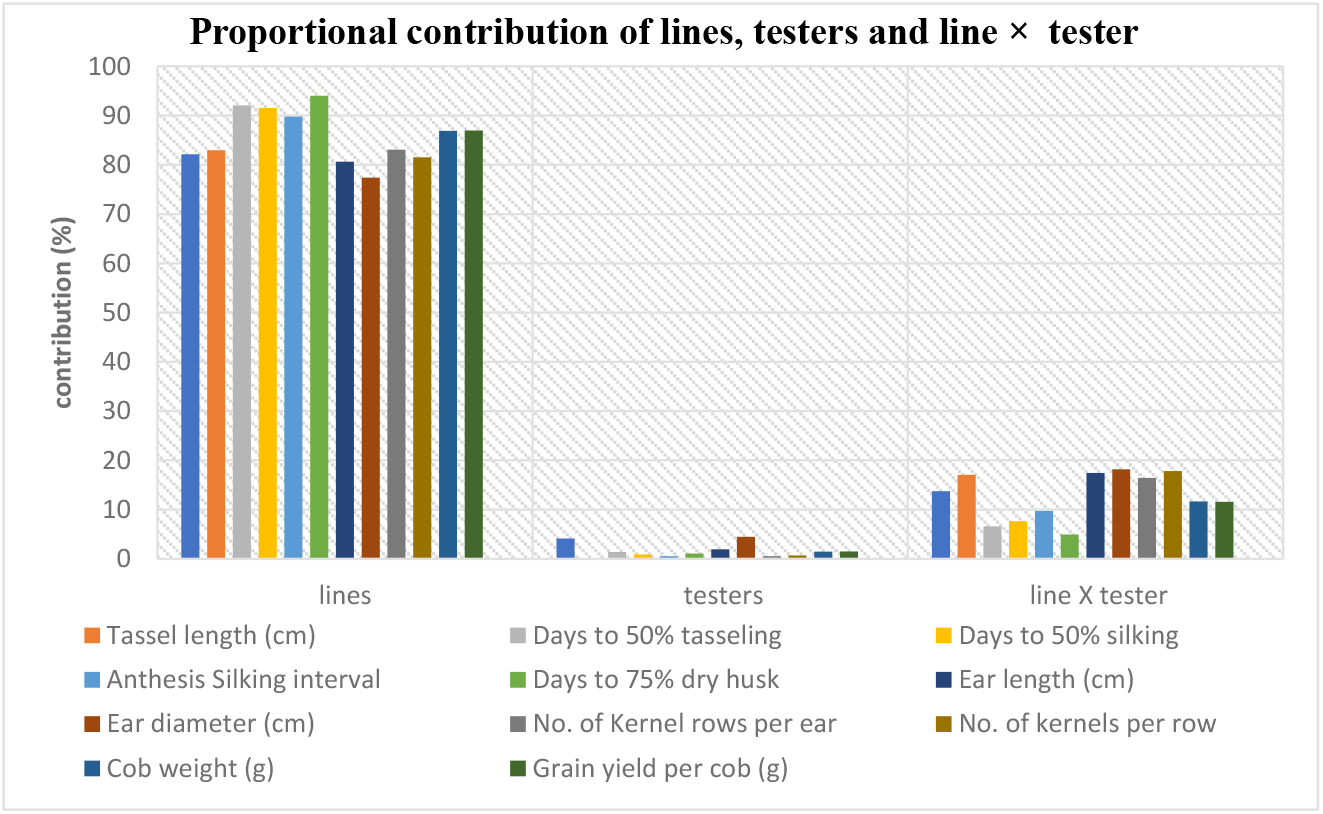
Proportional contribution of lines, testers, and line × tester interactions to total variance for all the traits under study.

### Molecular analysis

58 hybrids were developed in *Kharif*, 2022, and were evaluated in *Rabi*, 2022-23. Of the 58 female lines corresponding to 58 hybrids, 38 female lines were crossed with WS-5, 4 lines were crossed with WS-1, and 8 lines were commonly crossed to both wild species; this sums to a total of 50 female lines. So, 50 lines along with two wild species were checked for parental polymorphism with 12 SSR markers. Among the 12 SSRs, only the *bnlg1185* marker amplified both wild species. So, this marker is employed for further study. *bnlg1185* showed polymorphism for 40 female lines out of 50 lines, so only these 40 lines were used for molecular analysis. Of the 40 lines, 28female lines were involved in a cross with only WS-5, 8 were involved in a cross with only WS-1, and only four female lines were commonly crossed with both testers. So, the 42 parents (lines and testers) and their 44 hybrids were further considered for molecular diversity analysis. The banding pattern of all the hybrids showed both the amplicons present in the female as well as in the male parent, thus confirming the genuine crossing and heterozygotic condition of the hybrid. The amplification pattern and gel images of parents along with hybrids were displayed in Figures 4 and 5. Gel images developed were used for scoring, and the scored data were utilised for grouping of hybrids and parents using the Unweighted Pair Group Method with Arithmetic Mean (UPGMA) method in Darwin software version 6.0.021 (Perrier and Jacquemoud, 2006).

**Figure 4.**
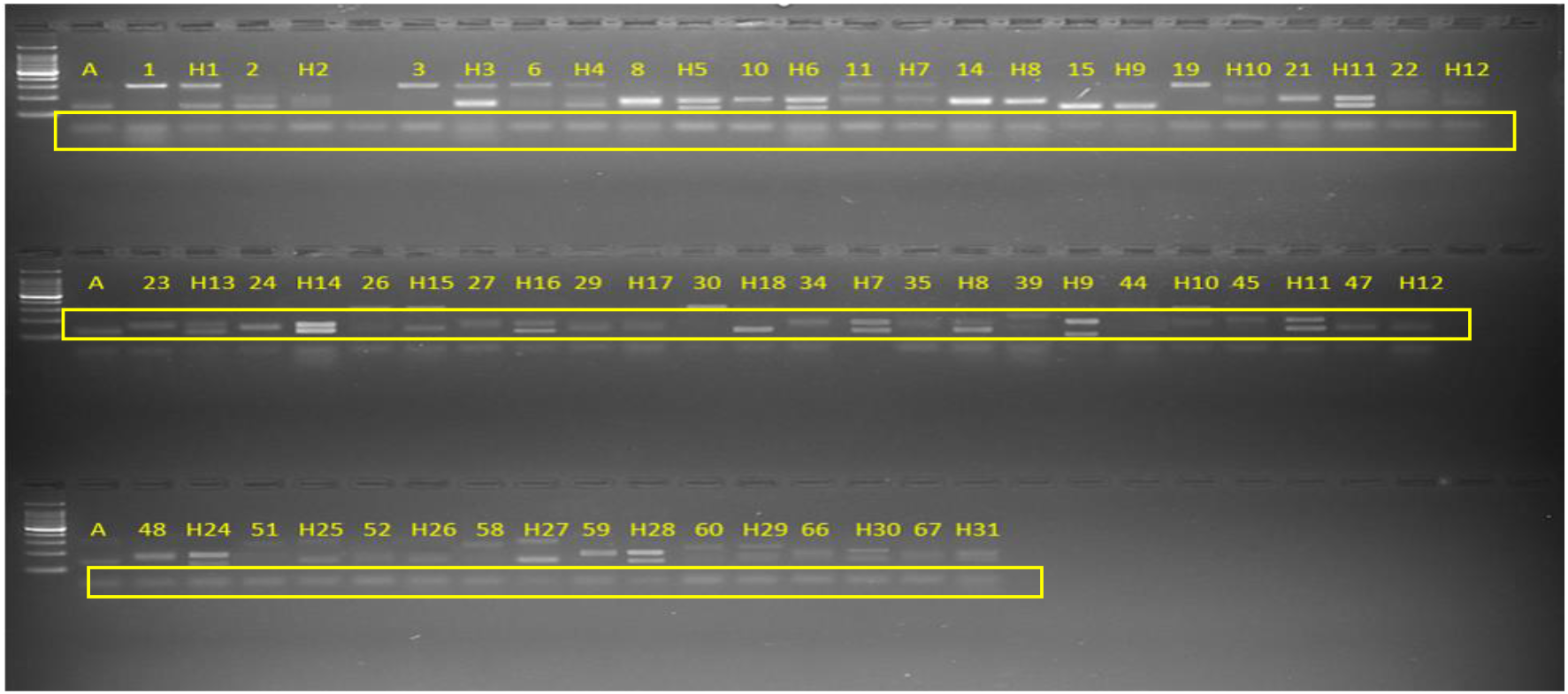
Molecular profile of parents along with their respective hybrids for hybrids developed from WS-5.

**Figure 5.**
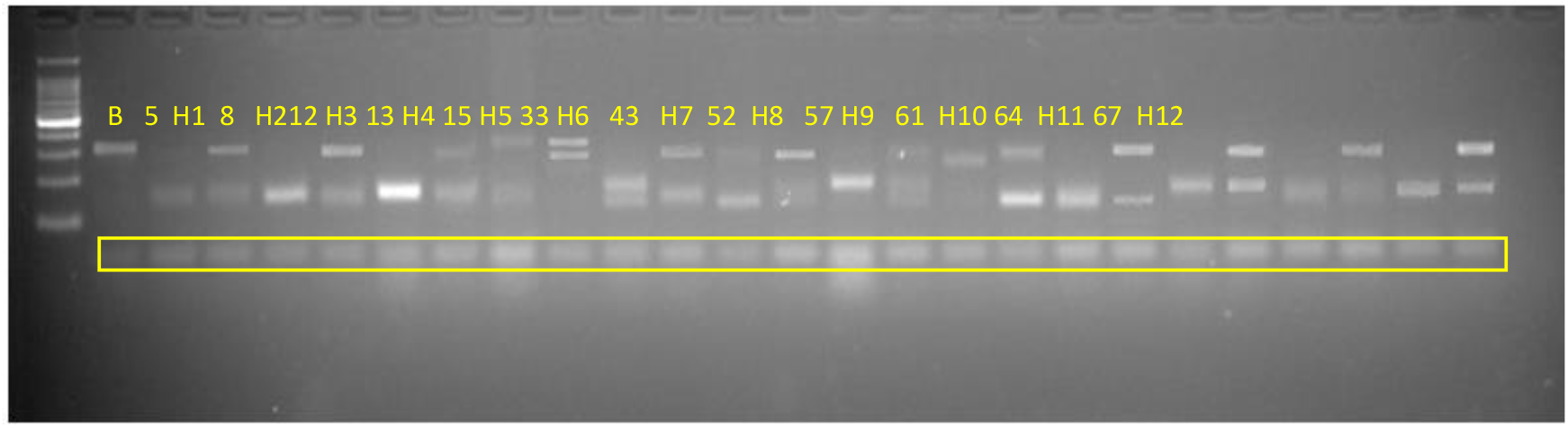
Molecular profile of parents along with their respective hybrids for hybrids developed from WS-1. **Where**, **A represents WS-5**, **B represents WS-1**, **Numbers represent female lines**, **H1, H2…, etc represent hybrids**

The UPGMA cluster analysis (Figure 6) categorized a total of 44 hybrids and 42 parent lines into two distinct groups, *viz*., Group A and Group B. Group A is the smallest group, in which WS-1 tester was grouped along with 12 hybrids and eight inbred lines, and all the 12 hybrids were developed from WS-1. Group B is the largest group, featuring the WS-5 tester along with 32 hybrids and 32 inbred lines. Within Group B, all 32 hybrids were developed from the WS-5 parent. Additionally, 32 inbred lines within Group B were further subdivided into two subgroups based on their distinct amplification patterns. All the hybrids showed both the amplicons present in the female as well as the male parent, thus confirming the genuine crossing and heterozygotic condition of the hybrid. These results were in accordance with Ahmed *et al*. (2019), Kumar *et al*. (2019).

**Figure 6.**
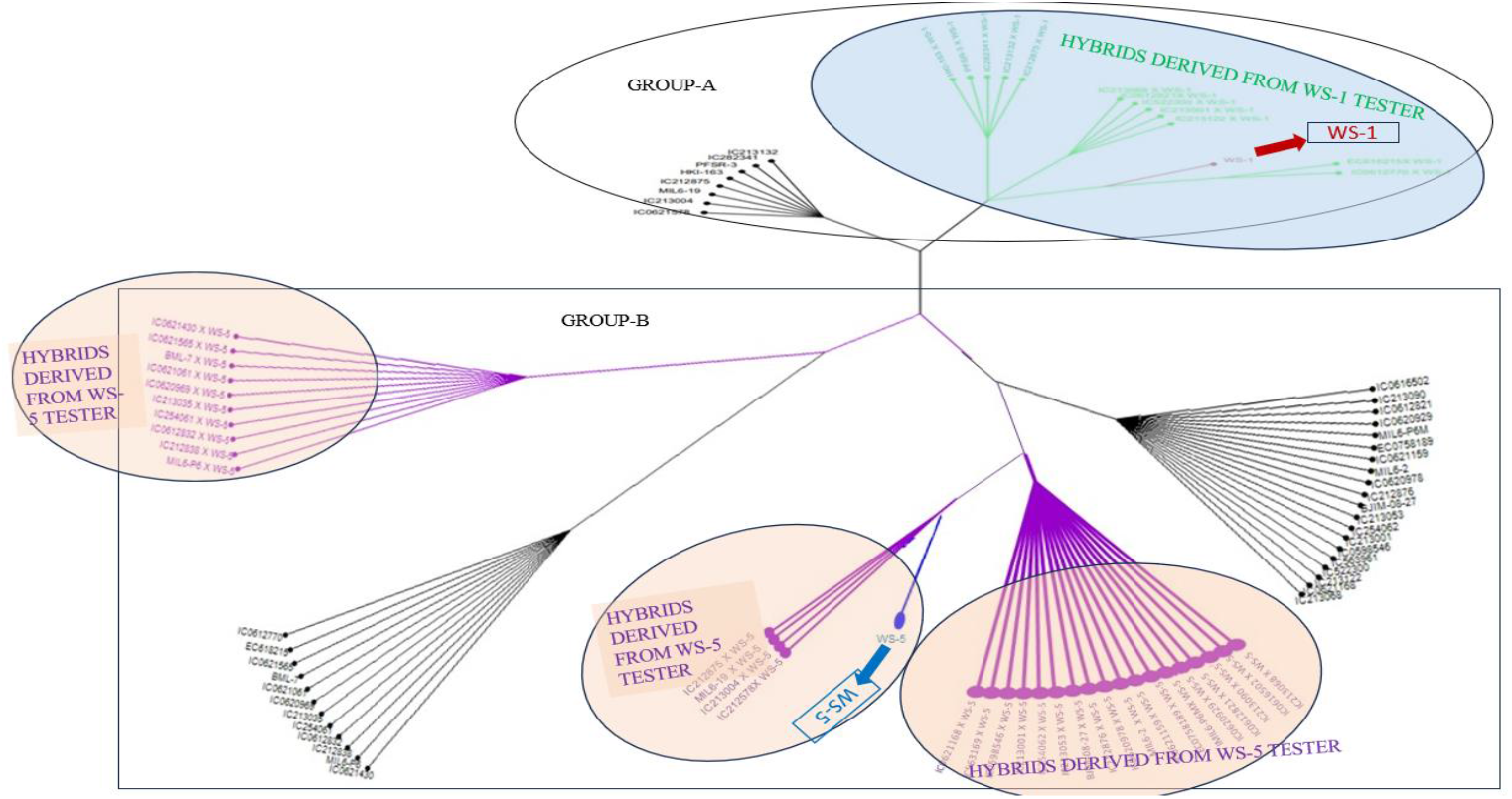
Clustering pattern of parents along with hybrids constructed by using Darwin software version 6.0.021.

## CONCLUSION

Introduction of wild species in the breeding program helps in transferring novel genetic variation into cultivated elite genotypes, improves genetic diversity and valuable traits, various biotic and abiotic stress resistance, to enhance heterosis and conserve genetic resources. So, the use of wild species helps in crop improvement. In the present research programme, inbred lines were crossed with wild species, WS-5 and WS-1. It was observed that hybrids showed typical wild characters such as tillering and prolificacy on the main tiller and side tillers. Prolificacy on the main tiller and side tillers, which was absent in modern cultivars. This multiple tillering habit was very useful for fodder purposes. GCA effects revealed that tester, WS-1, was a good general combiner for most of the traits and lines, MIL6-P6M, MIL6-26, MIL6-19, MIL6-1, IC213007, IC0621565, and IC0621049 were effective for ear traits and grain yield. Whereas the SCA effects showed that crosses, PFSR-3 × WS-5, EC618215 × WS-1, and IC213122 × WS-1 were good specific combiners for economically important traits. In this study, it was observed that significant SCA effects resulted from crosses between high × low, low × high, low × low, and high × high GCA combinations. Therefore, from the present study, it was learnt that to obtain a high SCA effect, we can include one low general combiner along with one high general combiner in future hybridization programs. Molecular analysis was carried out to determine the genuine nature of hybrids, and it was found that all hybrids were heterozygotic. Cluster analysis showed clear gene introgression from wild species in the hybrids, as hybrids grouped with their respective wild species. Most inbred lines clustered with WS-5, indicating greater compatibility with WS-5 than WS-1, which likely led to more successful crosses with WS-5. Whereas WS-1 was distantly related to most of the inbred lines, this indicates the presence of some crossing barriers with the inbred lines. Therefore, most of the crosses were not successful with the WA-1 wild species. This study revealed that the introduction of wild species will help in the creation of novel variation in the modern-day cultivars, as hybrids possess typical wild characteristics like tillering and prolificacy on main tillers and side tillers, which were absent in modern-day cultivars. Tillering habitat is economically useful for fodder purposes.

